# Identification of a master regulator of differentiation in *Toxoplasma*

**DOI:** 10.1101/660753

**Authors:** Benjamin S. Waldman, Dominic Schwarz, Marc H. Wadsworth, Jeroen P. Saeij, Alex K. Shalek, Sebastian Lourido

## Abstract

*Toxoplasma gondii* chronically infects a quarter of the world’s population, and its recrudescence can cause life-threatening disease in immunocompromised individuals and recurrent ocular lesions in the immunocompetent. Chronic stages are established by differentiation of rapidly replicating tachyzoites into slow-growing bradyzoites, which form intracellular cysts resistant to immune clearance and existing therapies. Despite its central role in infection, the molecular basis of chronic differentiation is not understood. Through Cas9-mediated genetic screening and single-cell transcriptional profiling, we identify and characterize a putative transcription factor (BFD1) as necessary and sufficient for differentiation. Translation of BFD1 appears to be stress regulated, and its constitutive expression elicits differentiation in the absence of stress. As a Myb-like factor, BFD1 provides a counterpoint to the ApiAP2 factors which dominate our current view of parasite gene regulation. Overall, BFD1 provides a genetic switch to study and control *Toxoplasma* differentiation, and will inform prevention and treatment of chronic infection.

## INTRODUCTION

The duration of infection is a critical parameter in the evolutionary fitness of infectious organisms. Pathogens can extend the period of infection by establishing a latent or chronic state, avoiding clearance through slow replication, altered immunogenicity, and a diminished impact on the host. These reservoirs are frequently resistant to treatment due to decreased metabolic rates. Such persistent stages can recrudesce or contribute to transmission, and are important barriers to curing and eradicating infectious diseases.

Within the phylum Apicomplexa, chronic stages play important roles in the life cycles of many of these pathogens. *Plasmodium vivax* hypnozoites in the liver are resistant to most antimalarial therapies, leading to long periods of latency followed by recurrence, complicating eradication efforts (Baird, 2009). *Toxoplasma gondii* tachyzoites are capable of invading any nucleated cell of warm-blooded animals, disseminating throughout the body and causing pathology through lysis of host cells. A proportion of tachyzoites differentiate into slow-growing bradyzoites, forming intracellular cysts with a tropism for brain and muscle tissue (Dubey et al., 1998). These cysts cannot be cleared by the immune system or by current therapies, and as a result up to a quarter of the world’s population is chronically infected with *Toxoplasma* (Montoya and Liesenfeld, 2004). *Toxoplasma* infection is life-threatening in immunocompromised individuals, and a majority of these cases result from recrudescent infections (Porter and Sande, 1992). Approximately 2% of infections result in ocular lesions—a leading cause of infectious blindness—with high rates of reactivation from chronic stages that persist after treatment (Jones et al., 2015).

Major changes accompany differentiation of rapidly proliferating tachyzoites into cyst-forming bradyzoites. The parasitophorous vacuole that *Toxoplasma* replicates within is modified into a heavily glycosylated cyst wall, containing many stage-specific proteins of unknown function (Coppin et al., 2005; Ferguson, 2004; Tomita et al., 2013; 2017). Parasite metabolism changes drastically, relying on anaerobic glycolysis instead of aerobic respiration and accumulating cytoplasmic starch granules (Denton et al., 1996; Guérardel et al., 2005; Sugi et al., 2017). Underpinning these dramatic changes in lifestyle, different studies have found hundred to thousands of genes are differentially regulated between tachyzoites and bradyzoites derived *in vitro* or *in vivo* (Behnke et al., 2008; Buchholz et al., 2011; Cleary et al., 2002; Lescault et al., 2010; Manger et al., 1998; Naguleswaran et al., 2010; Pittman et al., 2014; Radke et al., 2005; Ramakrishnan et al., 2019; Yahiaoui et al., 1999). Although differentiation can be induced *in vitro* through a variety of methods—including alkaline pH, heat shock, small molecules, and nutrient starvation—the molecular mechanisms driving bradyzoite differentiation remain poorly understood (Donald et al., 2002; Fox et al., 2004; Radke et al., 2006; Soete et al., 1994).

Attempts to identify mutants unable to differentiate have yielded strains with decreased rates of stage conversion, yet linking these phenotypes to inactivation of individual genes has proven challenging (Matrajt et al., 2002; Singh et al., 2002). A single validated class of apicomplexan transcription factors, the AP2 DNA-binding proteins (ApiAP2s), has been extensively investigated as potential regulators of differentiation. Knockouts of individual ApiAP2s modulate, but ultimately fail to completely ablate, bradyzoite differentiation, leading to the model that no master transcriptional regulator of this process exists in *Toxoplasma* (Hong et al., 2017; Huang et al., 2017; Jeffers et al., 2018; Radke et al., 2013; 2018; Walker et al., 2013). Overturning this view, we describe the identification and characterization of a transcription factor, Bradyzoite-Formation Deficient 1 (BFD1), that is both necessary and sufficient for differentiation. Bearing homology to the Myb transcription factor family, BFD1 represents a new paradigm for gene regulation in apicomplexans. The discovery also provides a unique molecular handle to study the chronic stages of infection, which represent a major barrier for developing radical cures against *Toxoplasma* and live-attenuated vaccines.

## RESULTS

### Generation of a differentiation reporter compatible with Cas9-mediated screens

To screen for mutants deficient in differentiation, we adapted the ME49 strain of *Toxoplasma* to make it compatible with Cas9-mediated gene disruption and enrichment of differentiated parasites (Sidik et al., 2014; 2016). Our reporter strain constitutively expresses RFP, and conditionally expresses the bright green fluorescent protein mNeonGreen (mNG) under the promoter of the canonical bradyzoite-specific gene *BAG1* (Fig. 1A). Our reporter strain further expresses Cas9, and selection for a guide RNA (gRNA) targeting the major tachyzoite surface antigen SAG1 resulted in 98% SAG1^−^ parasites, confirming robust inactivation of genes in this background (Fig. 1B). Growth of the reporter strain under alkaline stress, which induces *Toxoplasma* differentiation *in vitro*, resulted in increasing proportions of parasites expressing mNG (Fig. 1C, **S1A**).

**Figure 1.**
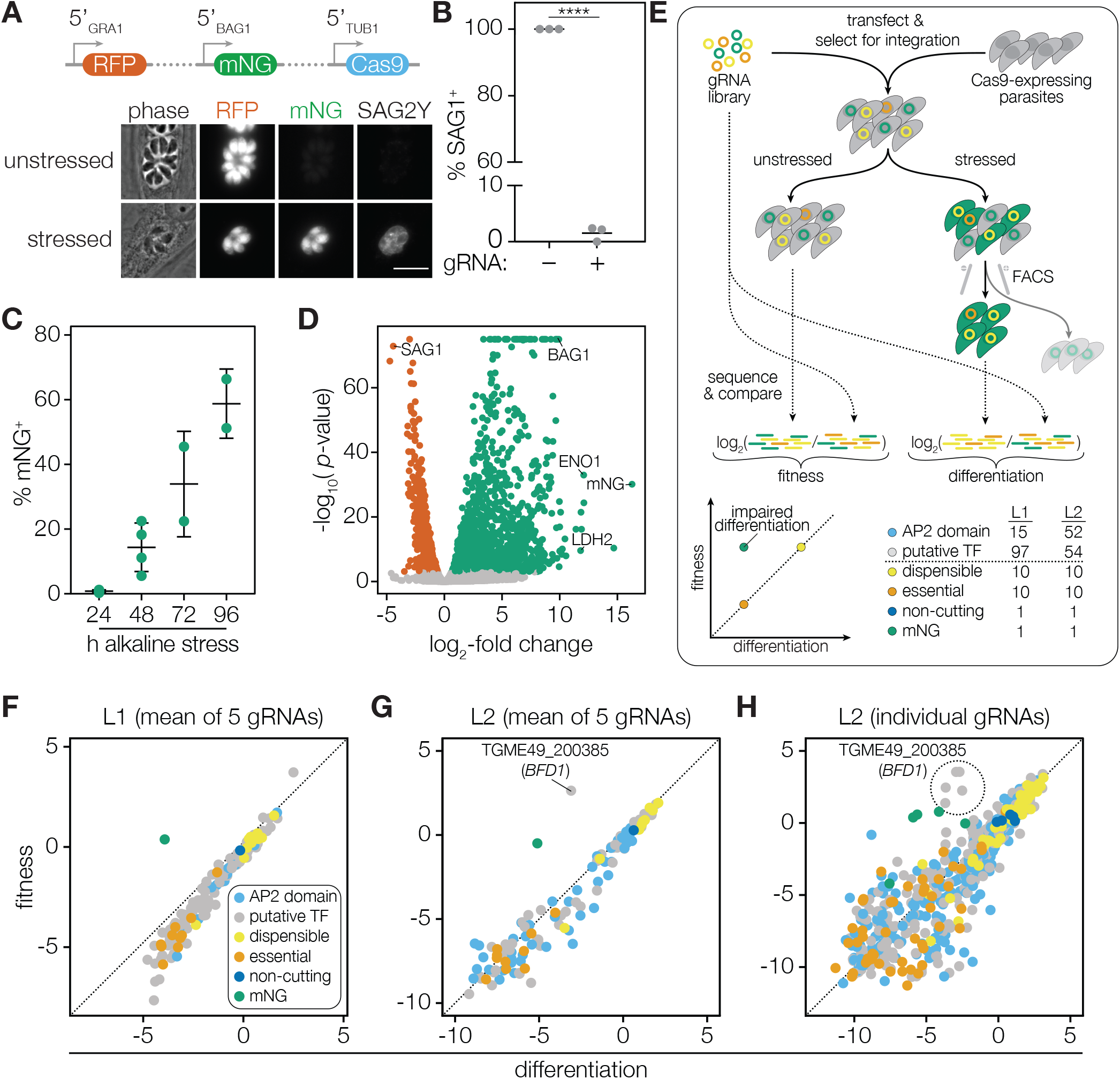
A genetic screen identifies a putative regulator of *T. gondii* differentiation. (**A**) Construction of a differentiation reporter strain that constitutively expresses RFP and Cas9, and conditionally expresses mNeonGreen (mNG) under the regulation of the bradyzoite-specific *BAG1* promoter, SAG2Y is a bradyzoite-specific surface marker. Images were taken after 48 h of growth under unstressed or alkaline stressed conditions. Scale bar is 10 µm. (**B**) Transfection and selection for a gRNA targeting the surface antigen SAG1 resulted in gene disruption in 98% of the resulting population. *n* = 3 biological replicates. 92–102 vacuoles were scored for each replicate. **** *p*-value < 0.0001; Student’s two-tailed *t*-test. (**C**) Percent of alkaline stressed reporter parasites expressing mNG, quantified by FACS. *n* = 2–4 biological replicates. Mean ± SD plotted. (**D**) RNA-sequencing and differential expression (DE) analysis identified 1,311 genes as significantly upregulated and 933 genes as significantly downregulated in bradyzoites (adjusted *p* < 0.001). *n* = 3 independent experiments. (**E**) Screening and analysis workflow. The log_2_-fold changes from the input library to the final unstressed or stressed mNG^+^ timepoints are defined as fitness or differentiation scores, respectively. (**F–G**) Fitness and differentiation scores at the gene level following screening L1 (F) or L2 (G). (**H**) Fitness and differentiation scores for individual gRNAs in L2.

To characterize transcriptomic differences between tachyzoites and bradyzoites, we performed stage-specific bulk RNA-sequencing using our reporter strain. We compared gene expression of FACS-purified tachyzoites (mNG^−^, 24 h unstressed growth) to bradyzoites (mNG^+^, 48 h stressed growth), and identified 1,311 genes as upregulated and 933 genes as downregulated in bradyzoites (Fig. 1D, **Data S1, Materials and Methods**). This comparison was chosen based on reproducible differences observed between 24 and 48 h unstressed replicates, which likely represent increased numbers of extracellular parasites following the completion of the lytic cycle at 48 h. A principal component analysis showed that 98% of variance is explained by growth condition, with minimal batch effects (**Fig. S1B**). Highly regulated genes agreed with previous datasets, with the canonical bradyzoite-specific genes *BAG1*, *LDH2*, and *ENO1* strongly upregulated, and the major tachyzoite surface antigen *SAG1* strongly downregulated. (Fig. 1D, **S1C**). Genes that were not previously annotated as differentially regulated tend to have lower expression, suggesting enhanced sensitivity in our dataset (**Fig. S1D**).

### Genetic screening identifies a putative regulator of *T. gondii* differentiation

Based on our previous CRISPR-based screens in the lab-adapted RH strain of *Toxoplasma*, we developed a screen for differentiation in a strain that retains normal stage conversion (ME49), and included a series of controls to benchmark the screening results (Sidik et al., 2016). However, executing these screens in a non-lab-adapted strain presented additional challenges. In particular, the lower viability and integration rates we observed limit the number of genes that can be screened simultaneously to the low hundreds (**Fig. S2**). By combining differential expression analysis with domain annotation and gene ontology, we assembled two libraries, each targeting ~100 potential nucleic acid–binding proteins, with 5 gRNAs per gene. Library 1 (L1) largely consists of genes identified as differentially regulated in our preliminary RNA-seq experiment. Library 2 (L2) contains genes with DNA-binding domains commonly found in transcription factors, such as zinc finger and Myb-like domains. Across both libraries, all 67 members of the ApiAP2 transcription factor family are targeted, along with 151 putative nucleic acid-binding proteins (Fig. 1E, **Data S2**) (Balaji et al., 2005; Painter et al., 2011). As controls, each library additionally contains 10 genes known to be essential, 10 genes known to be dispensable, 10 non-cutting gRNAs, and 5 gRNAs against the mNG reporter itself.

Following transfection of the libraries, parasites were maintained in selective medium for four passages to allow for integration of the gRNA plasmids before splitting the population between unstressed or alkaline-stressed conditions for 10 days (Fig. 1E, **Materials and Methods**). After the 10 days, mNG^+^ parasites from the stressed populations were isolated by FACS. Integrated gRNAs from the final unstressed population or from mNG^+^ stressed population were amplified and sequenced along with the input library. The fold-change in relative abundance between each final sample and the library was calculated for each gRNA. The mean log_2_ fold-change for guides against each gene are referred to as a fitness or differentiation score, based on comparisons to the unstressed or mNG^+^ stressed samples, respectively. Candidate genes should be depleted specifically in the mNG^+^ population (low differentiation score relative to their fitness score), as should gRNAs against the mNG reporter. In L1, only control mNG gRNAs had low differentiation scores compared to their fitness scores (Fig. 1F). In L2, however, gRNAs targeting a single gene—TGME49_200385, which we name Bradyzoite-Formation Deficient 1 (*BFD1*)—were specifically depleted in the differentiated populations along with control mNG gRNAs (Fig. 1G–H). Failure of a *BFD1* mutant to express the mNG reporter following alkaline stress was confirmed by transfecting a single *BFD1* targeting gRNA into our reporter strain. A frameshifted clone was isolated with a single nucleotide insertion at the cut site (*BFD1*^frameshift^, **Fig. S3**). Under alkaline stress, the wild-type reporter strain showed robust mNG expression, while the *BFD1*^frameshift^ clone did not (**Movie S1**).

### BFD1 contains conserved DNA-binding domains and localizes to the nucleus

We defined the sequence of the *BFD1* open reading frame based on cDNA sequencing, which differed from the annotated gene model and encoded a protein of 2,415 amino acids (**Fig. S4**). BFD1 contains two tandem SANT/Myb-like DNA-binding domains (SMART accession 00717), flanked by large extensions lacking identifiable motifs. Phylogenetic analysis of the first and second domain revealed homology to the R2 and R3 repeats of the human c-Myb protein, respectively (Fig. 2A–B, **Fig. S5**). *BFD1* orthologs can be found in all apicomplexan species that form tissue cysts, while different patterns of conservation are found for other SANT-domain–containing proteins. As might be expected for a putative transcription factor, transiently expressed C-terminally epitope-tagged BFD1 showed nuclear localization (Fig. 2C).

**Figure 2.**
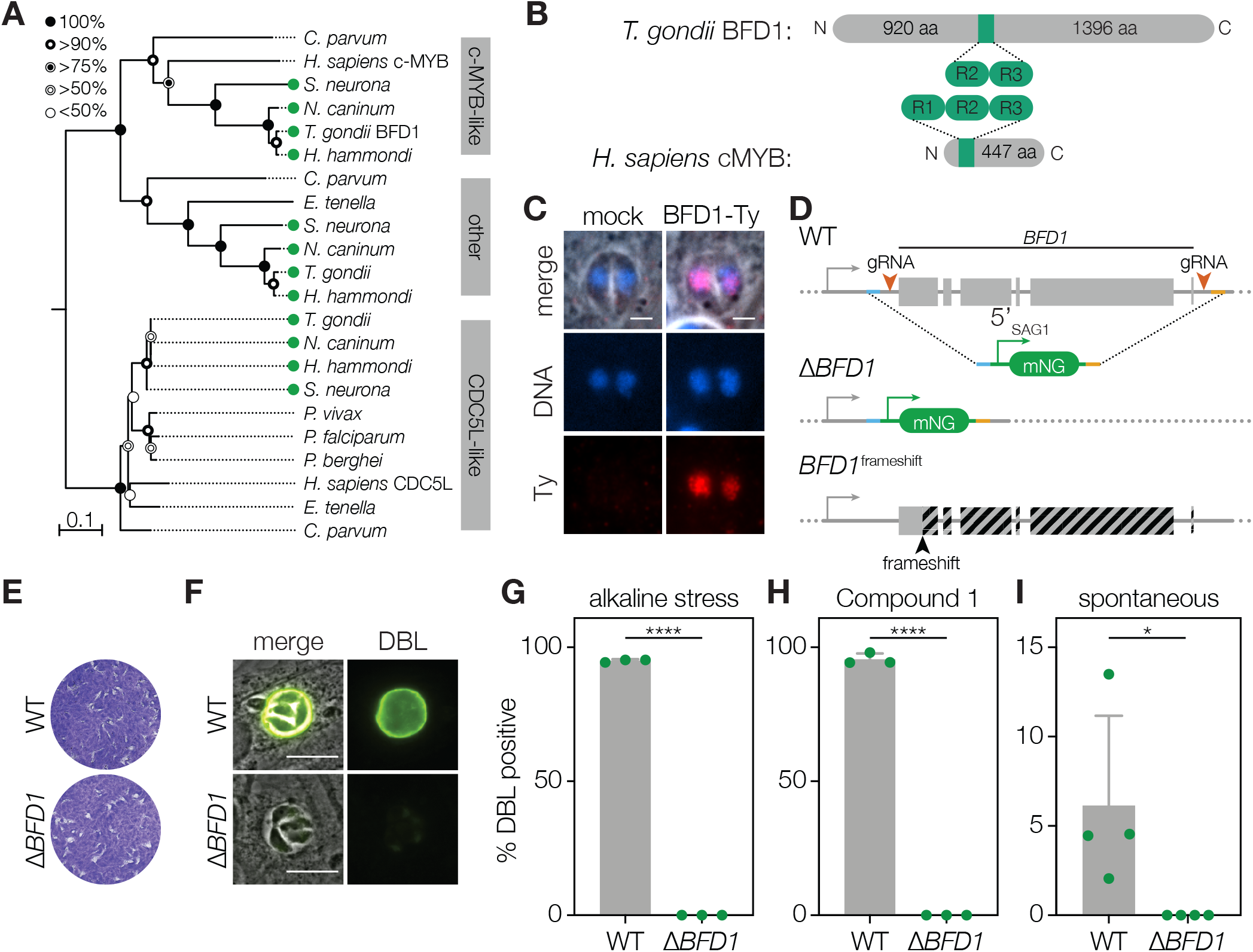
BFD1 is a nuclear factor necessary for differentiation. (**A**) Neighbor-joining tree showing the phylogenetic relationship of the concatenated SANT domains from BFD1 and its closest homologs in other apicomplexans and humans. BFD1 forms a clade with human c-Myb, distinct from CDC5L-like sequences. Tissue-cyst forming species indicated (green circle). Bootstrap values for 1000 trials are displayed. Scale bar is substitutions per site. (**B**) Diagram of BFD1 and human c-Myb highlighting the SANT domains (green). The DNA-binding repeats of BFD1 are similar to the second and third repeats of c-Myb. (**C**) Wild-type parasites transfected with a C-terminally Ty-tagged *BFD1* cDNA, driven by the *TUB1* promoter, show nuclear localization of the transgene. DNA stained with Hoechst (blue) and Ty immunostained with BB2 (red). Scale bar is 3 µm. (**D**) Generation of a *BFD1* knockout through homologous recombination in ME49∆*KU80*. The entire coding sequence of *BFD1* was replaced with an mNG-expression cassette using gRNAs against the 5′ and 3′ ends. Diagram of frameshift mutant in the reporter background (*BFD1*^frameshift^) shown for comparison. (**E**) Plaque assays of parental ME49∆*KU80* (WT) or ME49∆*KU80*∆*BFD1* (∆*BFD1*) grown under unstressed conditions for 14 days. (**F**) Representative vacuoles of wild-type or ∆*BFD1* parasites after 48 h of alkaline stress. FITC-labeled *Dolichos biflorus* lectin (DBL) specifically stains differentiated vacuoles. Scale bar is 10 µm. (**G–I**) Quantification of differentiation in WT and ∆*BFD1* parasites following 48 hours of alkaline stress (G), 48 hours of Compound 1 treatment (H) or occurring spontaneously (I). Mean ± SD of *n =* 3–4 biological replicates. % DBL positive vacuoles calculated from at least 100 vacuoles counted for each replicate. **** *p*-value < 0.0001, * *p*-value < 0.05, Student’s one-tailed *t*-test.

### Loss of BFD1 blocks parasite differentiation regardless of induction method

To provide a clean background for precise genetic manipulation, we generated a low-passage, NHEJ-deficient ME49 strain through deletion of *KU80* (**Fig. S6**). In this background, we replaced the entire coding sequence of *BFD1* with an mNG expression cassette (Fig. 2D). Deletion of *BFD1* caused no defect in tachyzoite growth as assayed by plaque formation (Fig. 2E). Differentiating vacuoles can be distinguished using *Dolichos biflorus* lectin (DBL) staining, which recognizes *N*-acetylgalactosamine on the bradyzoite-specific cyst-wall protein CST1 (TGME49_264660) (Tomita et al., 2013). Wild-type vacuoles became robustly DBL^+^ after 48 h under alkaline stress. By contrast, no ∆*BFD1* vacuoles developed DBL positivity under identical conditions (Fig. 2F–G). ∆*BFD1* parasites exhibited a similar defect when differentiation was induced by the small molecule Compound 1 or occurred spontaneously (Fig. 2H–I) (Donald et al., 2002; Radke et al., 2006).

### Characterizing *Toxoplasma* cell-cycle progression and differentiation by single-cell RNA-sequencing

To profile cell-cycle progression and the asynchronous process of differentiation, we performed single-cell RNA-sequencing (scRNA-Seq) of *T. gondii* using Seq-Well (Gierahn et al., 2017). Wild-type or ∆*BFD1* parasites were grown under unstressed or stressed conditions (24, 48 or 72 h), mechanically released from host cells, and analyzed. Following downstream processing and alignment, 26,560 cells passed quality control cutoffs, with on average 1,537 unique molecular identifiers (UMIs, a proxy for unique transcripts) and 685 genes represented per cell (**Fig. S7, Materials and Methods**). As cells from the 72 h timepoint were of the highest quality, we clustered unstressed parasites from this timepoint to examine the tachyzoite cell cycle. Seven clusters were identified, with six arranged in a circular pattern in a uniform manifold approximation and projection (UMAP) visualization (Fig. 3A) (McInnes et al., 2018). Scoring cells based on expression of known cell-cycle signatures identified clusters 0 and 1 as G1-like and clusters 2, 3, 5 and 6 as S/M-like (Fig. 3B, **Data S3**) (Behnke et al., 2010; Suvorova et al., 2013). We identified 1,173 genes as upregulated in at least one cluster compared to all the others, including 525 genes not previously identified as cell-cycle regulated. Plotting the average expression profile of these markers across an existing dataset of synchronized tachyzoite gene expression revealed a progression through the cell cycle in a counter clock-wise direction around the plot (Fig. 3A, 3C, **Data S4**) (Behnke et al., 2010). The proportion of cells identified as being in G1 or S/M matches the 60 to 40 ratio previously measured (Fig. 3D) (Radke et al., 2001).

**Figure 3.**
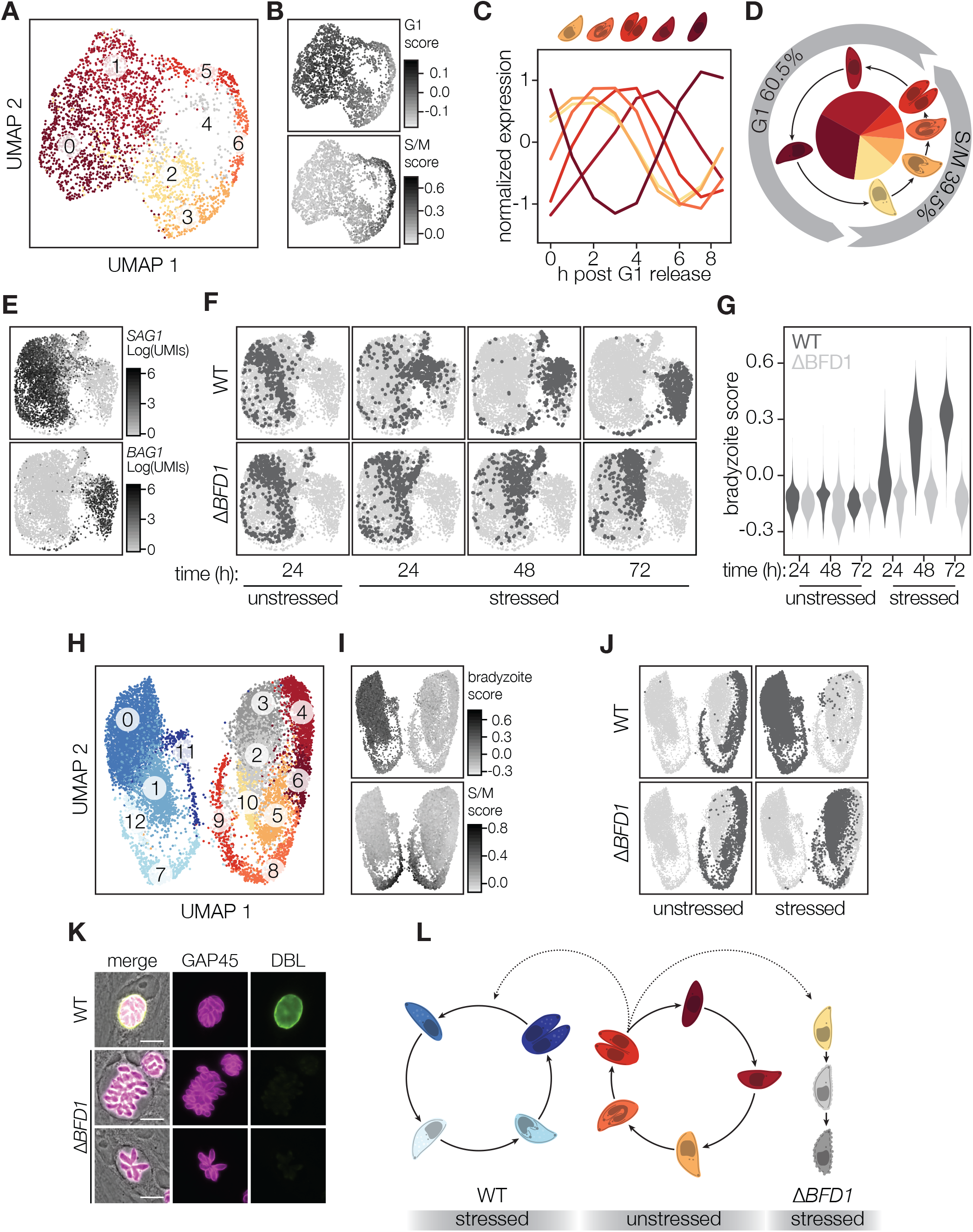
BFD1 is required for initiation of differentiation and expression of bradyzoite genes. (**A**) Clustering of unstressed parasites of both genotypes after 72 h of growth (3,149 cells total). Visualized by uniform manifold approximation and projection (UMAP). (**B**) Scoring of cells based on expression of known G1 or S/M-specific gene sets. (**C**) Expression profiles for genes identified as differentially expressed by cells within each cluster in A compared to cells within all other clusters, derived from a microarray dataset of synchronized tachyzoites (Behnke et al., 2010). (**D**) Proportion of cells in G1 or S/M clusters. (**E**) Cells colored by expression of canonical stage-specific genes *SAG1* and *BAG1* following clustering of all parasites from all timepoints, genotypes and growth conditions. UMAP visualization is downsampled to 500 cells from each combination of timepoint, genotype and growth condition (6,000 cells total). (**F**) UMAP as in E, separated by sample of origin. (**G**) Scoring cells by sample of origin based on expression of genes highly upregulated in bradyzoites. (**H**) Visualization by UMAP of clustering of all WT and ∆*BFD1* parasites from unstressed and stressed cultures at 72 h timepoint. (**I**) UMAP projection as in H, scored by expression of highly upregulated bradyzoite genes as in G or S/M-specific genes as in B. (**J**) UMAP as in H, separated by sample of origin. (**K**) Representative WT and ∆*BFD1* vacuoles at 72 h post alkaline stress. GAP45 is a marker for the inner membrane complex. Scale bar is 10 µm. (**L**) Proposed model of cell cycle progression for WT and ∆*BFD1* parasites under the different treatments.

Clustering cells from all timepoints, growth conditions and genotypes revealed a clear division between tachyzoite (SAG1^+^) and bradyzoite (BAG1^+^) clusters (Fig. 3E). Under alkaline stress, wild-type parasites quickly exit the tachyzoite cell cycle and begin progressing towards bradyzoite-containing clusters (Fig. 3F). Scoring cells based on their expression of the 100 most upregulated genes identified by our stage-specific RNA-sequencing shows a clear increase in this signature over time (Fig. 3G, **Data S3**). Consistent with the robust DBL positivity seen at 48 h post induction, *CST1* is among the earliest genes induced (**Fig. S8A-B, Data S5**). Endogenous tagging of two additional strong, early markers of differentiation (TGME49_312330, TGME49_208740) identified both as cyst-wall proteins, confirming the localization recently shown for TGME49_208740 (**Fig. S8C**) (Tu et al., 2019). While *BFD1* transcription is not exclusive to bradyzoites, *BFD1* is identified as modestly upregulated at 72 h in stressed wild-type parasites by our scRNA-Seq data, and is significantly enriched in bradyzoite-containing clusters (**Fig. S9**, **Data S1, S5, S6**). UMAP visualization of unstressed or stressed parasites at the 72 h timepoint showed two circular patterns, driven primarily by cell-cycle and bradyzoite-specific gene signatures (Fig. 3H-I, **Fig. S10**). We identified 228 genes as upregulated specifically in replicating bradyzoites (clusters 7, 11 and 12), including the previously identified bradyzoite rhoptry protein 1 (**Fig. S11, Data S7**) (Schwarz et al., 2005). In wild-type parasites, 19% of bradyzoites are in clusters expressing S/M markers, recapitulating the approximately two-fold slower replication previously measured (Dzierszinski et al., 2004).

### BFD1 is necessary for bradyzoite differentiation

Under unstressed conditions, the distribution of ∆*BFD1* parasites mirrors that of wild-type parasites along the various tachyzoite clusters. However, tracking the fate of ∆*BFD1* parasites under stress conditions demonstrates that they continue to replicate as tachyzoites until the 72 h timepoint, when they cluster separately from both wildtype tachyzoites and bradyzoites (Fig. 3F). Scoring these cells based on their expression of bradyzoite-specific genes revealed a widespread failure of ∆*BFD1* parasites to initiate bradyzoite-specific gene expression (Fig. 3G, **Data S3, S5**). Under alkaline stress, while 7.2% of ∆*BFD1* parasites are clustered with dividing S/M phase tachyzoites, 88.6% of cells organize into three G1-adjacent clusters (2, 3, and 10) distinct from those harboring G1 tachyzoites (Fig. 3J). Aberrant morphologies observed for ∆*BFD1* parasites after 72 h under stress suggest that these G1-adjacent clusters contain dying parasites, although we cannot exclude the existence of a G1-arrested state (Fig. 3K). We propose that, due to the necessity of *BFD1* for differentiation, after 72 hours under alkaline stress, ∆*BFD1* parasites are dying due to their failure to respond appropriately to that stress (Fig. 3L).

### Overexpression of BFD1 causes differentiation in the absence of inducing conditions

Our stage-specific transcriptional analysis demonstrates that *BFD1* mRNA levels are constant across different stages, and present at a similar level to genes whose protein products are found in tachyzoites (Fig. 4A). We successfully introduced an epitope-tagged allele of *BFD1* into the endogenous locus of the ∆*BFD1* strain. Monitoring BFD1 levels by staining for the epitope tag revealed significant staining only under stress conditions (Fig. 4B). Inspection of the *BFD1* locus revealed an unusually long 5′ untranslated region (UTR), which at 2.7 kb in length falls in the 98^th^ percentile of all annotated 5′ UTRs in the *Toxoplasma* genome (Fig. 4C). The 5′ UTR of *BFD1* encodes four untranslated open reading frames (75–91 amino acids in length; Fig. 4D) that, like UTR length, characterize many post-transcriptionally regulated genes (Hinnebusch et al., 2016). To investigate whether overexpression of *BFD1* would influence differentiation in the absence of stress, we generated two constructs expressing epitope-tagged BFD1 under the *TUB1* promoter—either full-length (BFD1^WT^) or a mutant lacking the DNA-binding domains (BFD1^∆DBD^) (Fig. 4E). Transient overexpression of BFD1^WT^ induced differentiation in over 60% of wild-type or ∆*BFD1* parasites, demonstrating that BFD1 expression is sufficient to induce differentiation under unstressed conditions. Differentiation of ∆*BFD1* parasites transfected with the wild-type construct confirmed that their defect is specifically due to the deletion of *BFD1* (Fig. 4F). Overexpression of BFD1^∆DBD^ failed to induce differentiation in either genotype, demonstrating the requirement of the DNA-binding domains of BFD1 for activity and suggesting it acts as a transcription factor to drive differentiation.

**Figure 4.**
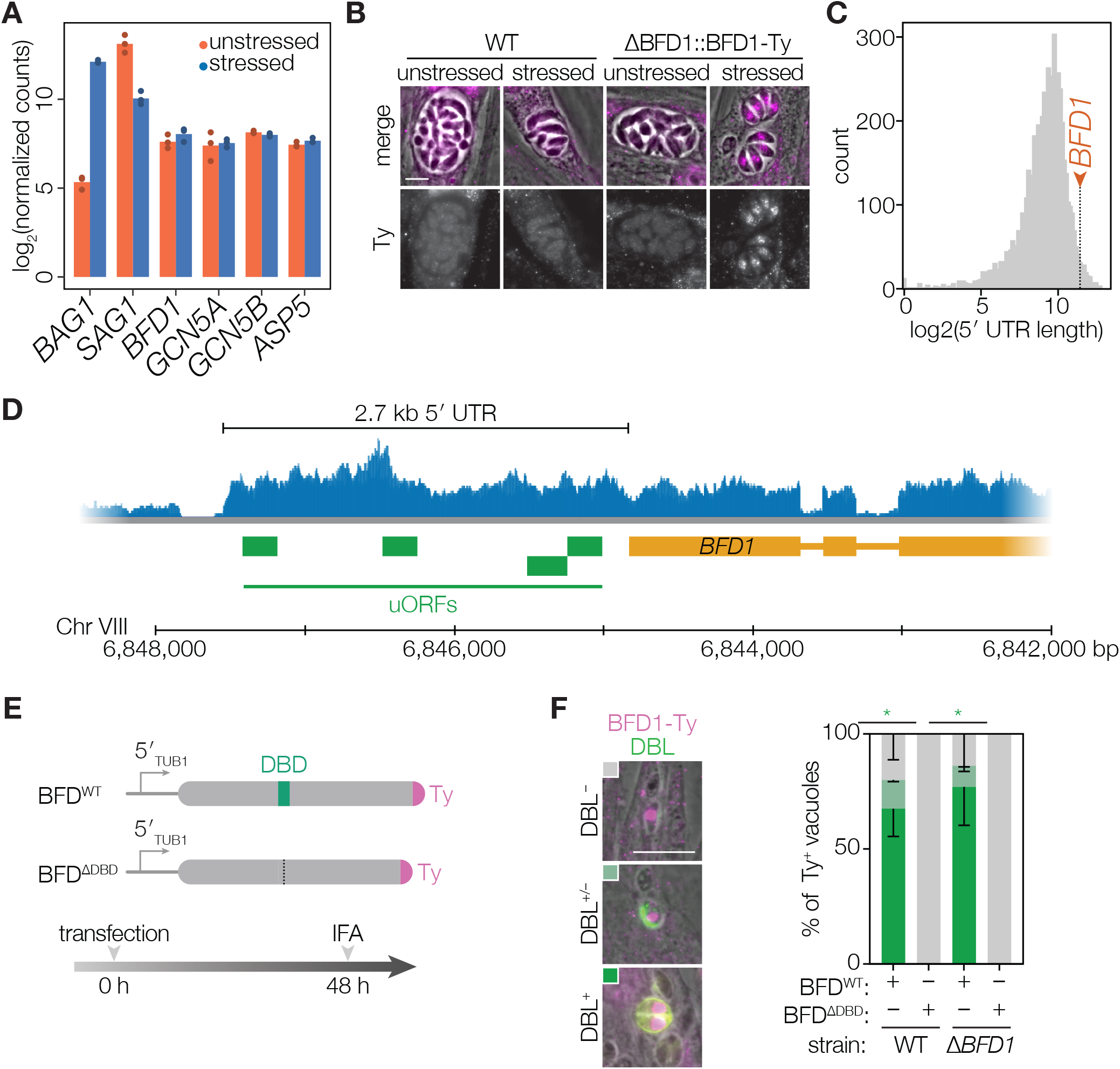
Overexpression of BFD1 is sufficient to induce differentiation in the absence of stress. (**A**) Normalized counts for selected genes in unstressed or stressed stage-specific RNA-sequencing replicates. (**B**) Representative WT and ∆*BFD1*::*BFD1*-Ty vacuoles after 48 h growth under unstressed or stressed conditions. Ty immunostained with BB2 (magenta). Scale bar is 5 µm. (**C**) Distribution of lengths of 5,699 previously annotated *Toxoplasma* 5′ UTRs. By manual annotation, the 5′ UTR of *BFD1* is 2,709 bp (red arrow), placing it in the 98^th^ percentile. (**D**) *BFD1* locus showing the RNA expression trace from tachyzoites (blue), position of predicted uORFs (green), and the main ORF (yellow). (**E**) Constructs and experimental workflow for transient overexpression of BFD1. Epitope-tagged cDNA copies of BFD1 (BFD1^WT^) or BFD1 lacking its DNA-binding domains (BFD1^∆DBD^) are under the regulation of the *TUB1* promoter. Parasites were immuno-labeled for Ty (magenta) and for differentiation with FITC-conjugated DBL (green). (**F**) Quantification of differentiation in WT or ∆*BFD1* parasites 48 h after transient overexpression of BFD1^WT^ or BFD1^∆DBD^. Ty^+^ vacuoles were identified and then scored for DBL positivity as shown in representative images. Scale bar is 10 µm. *n* = 2 independent replicates, 17–61 vacuoles counted per replicate. Mean ± SD plotted. * *p*-value < 0.05, Student’s two-tailed *t*-test.

## DISCUSSION

Differentiation from tachyzoites to bradyzoites establishes chronic *Toxoplasma* infection. However, the molecular pathways regulating this transition have remained unclear, despite evidence that disparate inputs—heat shock, alkaline stress, and nutrient starvation—converge on a common transcriptional program. Using bulk and single-cell RNA-sequencing, we characterize differentiation in unprecedented detail. Through Cas9-mediated genetic screens, we identify a single gene (*BFD1)* as indispensable for differentiation. ∆*BFD1* parasites grow normally under standard conditions but fail to differentiate under all induction conditions tested. Overexpression of BFD1 is sufficient to induce differentiation under standard conditions in both wild-type and knockout parasites, demonstrating its role as a master regulator of bradyzoite formation in *Toxoplasma*.

By profiling FACS-enriched differentiated populations, we captured transcriptional differences between tachyzoites and bradyzoites with greater sensitivity and dynamic range than achieved by previous datasets (Behnke et al., 2008; Buchholz et al., 2013; Lescault et al., 2010; Naguleswaran et al., 2010; Pittman et al., 2014; Radke et al., 2005; Ramakrishnan et al., 2019). These changes likely reflect a combination of factors including altered replication, nutrient availability, and general stress responses, in addition to the bradyzoite differentiation program. We note that 58% of genes identified as downregulated in our stage-specific RNA-sequencing are cell-cycle regulated, suggesting that changes in replication rate are responsible for many of the transcriptomic differences observed at the population level (**Data S1, S4**). The genetic control afforded by BFD1 on differentiation will help future studies disentangle the contributions of these variables.

A single family of DNA-binding proteins, the ApiAP2s, has been investigated for their role as *Toxoplasma* transcription factors and mediators of differentiation. While the phenotypes associated with many ApiAP2 mutants are striking, no single gene knockout has resulted in a complete block of differentiation, leading to the assumption that multiple transcription factors regulate *Toxoplasma* bradyzoite development (Hong et al., 2017; Huang et al., 2017; Jeffers et al., 2018; Radke et al., 2013; 2018; Walker et al., 2013). By screening a wider range of putative nucleic–acid binding proteins, especially those containing well-conserved DNA-binding motifs such as zinc finger and Myb-like domains, we discovered that inactivation of *BFD1* completely ablates bradyzoite formation. This does not preclude important roles for ApiAP2 proteins as downstream mediators of the differentiation program, and additional work is needed to identify the hierarchy of transcriptional regulation during induction. *Toxoplasma* encodes thirteen other proteins containing SANT/Myb-like domains, suggesting the existence of a second extensive transcription factor family. Myb domain–containing proteins are widespread among eukaryotes, and have been implicated in the regulation of encystation in *Entamoeba* and *Giardia*, along with a wide variety of stress responses in plants (Ambawat et al., 2013; Dubos et al., 2010; Ehrenkaufer et al., 2009; Huang et al., 2008; Sun et al., 2002). In humans, c-Myb is thought to function as a pioneer transcription factor, binding to chromatin and recruiting histone acetyltransferases to commit cells to specific hematopoietic lineages (Fuglerud et al., 2018; Sandberg et al., 2005). Among apicomplexan parasites, a Myb domain-containing protein has been identified as important for erythrocytic growth of *Plasmodium falciparum*, suggesting that other family members likely play important roles throughout the phylum (Boschet et al., 2004; Gissot et al., 2005).

ScRNA-Seq enables profiling of thousands of cells across asynchronous processes, and has been successfully used to examine commitment to sexual differentiation in *Plasmodium* spp. (Bancells et al., 2019; Poran et al., 2017). Implementing these approaches in *Toxoplasma* retained information about cell-cycle residency and the relative timing of gene expression that is lost in bulk analyses. Moreover, scRNA-Seq allowed us to identify novel markers specific to actively replicating bradyzoites and genes expressed during the earliest stages of differentiation (**Data S7**). This detailed view of differentiation revealed that *BFD1* knockout parasites progress normally throughout the tachyzoite cycle but fail to initiate bradyzoite differentiation following alkaline stress. *BFD1* therefore stands out from other genes known to influence differentiation for its complete essentiality for stage conversion.

As a necessary and sufficient regulator of differentiation, *BFD1* provides a focal point for the molecular mechanisms underlying differentiation. In *Plasmodium*, identification of AP2-G as the master transcriptional regulator of gametogenesis has permitted placement of multiple genes observed to affect sexual differentiation into a unified regulatory framework, and allowed directed investigation into their mechanisms of action (Brancucci et al., 2014; Coleman et al., 2014; Eksi et al., 2012; Filarsky et al., 2018; Ikadai et al., 2013; Kafsack et al., 2014; Sinha et al., 2014). The ability to induce synchronized sexual differentiation through conditional overexpression of AP2-G has allowed finer temporal mapping of the gene expression changes that accompany gametogenesis, and application of a similar approach in *Toxoplasma* is now possible using *BFD1* (Kent et al., 2018).

Identification of the genes regulated by BFD1, its binding sites, and its accessory factors will determine how it regulates differentiation, while studies into *BFD1* expression and post-translational regulation will help uncover its response to environmental stimuli. Our transcriptional profiling reveals that *BFD1* is expressed in the 67^th^ percentile in tachyzoites, and expression at this level or higher is corroborated by many other RNA-sequencing datasets (Buchholz et al., 2011; Hassan et al., 2017; Melo et al., 2013; Pittman et al., 2014; Ramakrishnan et al., 2019; Reid et al., 2012; Swierzy et al., 2017). *BFD1* expression therefore does not appear to be stage-specific despite a modest 1.5-to 3.6-fold upregulation in bradyzoites observed by bulk RNA-Seq or scRNA-Seq—the former below our cutoff for significance (**Data S1, S5**). However, expression of an epitope-tagged version of *BFD1* from the endogenous locus was only observed under stress conditions. In other eukaryotes, long 5′ UTRs containing small untranslated open reading frames—as observed for *BFD1*—are the locus of translational regulation (Hinnebusch et al., 2016). Taken together, these results suggest that regulation of BFD1 may be, at least in part, post-transcriptional. Preferential translation of some transcripts under stress conditions has been reported in *Toxoplasma*, and mutations in RNA-binding proteins have resulted in severe defects that suggest an important role for translational control during differentiation (Gissot et al., 2013; Narasimhan et al., 2008).

Future studies will need to address the function of *BFD1* during acute and chronic mouse infection. Mutations resulting in decreased rates of differentiation in cell culture generally display more profound defects in mice; however, the specificity of the ∆*BFD1* phenotype will help define the role of bradyzoites in pathogenesis and immunological memory during *Toxoplasma* infection (Al-Anouti et al., 2004; Huang et al., 2017; Jeffers et al., 2018; Saeij et al., 2008). ∆*BFD1* parasites may also represent an ideal attenuated vaccine strain, capable of proliferating robustly yet unable to enter a chronic state. Collectively, modulation of BFD1 holds substantial clinical and biotechnological potential, as chronic infection represents a major barrier to both the treatment of *Toxoplasma* and its use in delivery of heterologous antigens and protein-based therapeutics.

## MATERIALS & METHODS

### Strains and cell culture

Human foreskin fibroblasts were maintained in DMEM (Gibco) supplemented with 3% inactivated fetal serum (IFS) and 10 μg/mL gentamicin, referred to as standard medium. If HFFs were to be used in bradyzoite experiments, host cells were maintained exclusively in DMEM supplemented with 10% IFS and 10 μg/mL gentamicin prior to infection. Alkaline stress medium consists of RPMI 1640 (Sigma), supplemented with 1% IFS and 10 μg/mL gentamicin, and buffered with 50 mM HEPES adjusted to pH 8.1 with 10N NaOH. Compound 1 was used at 3 μM in standard medium (Donald et al., 2002; Radke et al., 2006).

### Plasmids and primers

Oligos were ordered from IDT. All cloning was performed with Q5 2× master mix (NEB) unless otherwise noted. Primers and plasmids used or generated in this study can be found in **Data S8**.

### Strain generation

#### Bradyzoite reporter strain

Starting with a robustly cyst-forming ME49 strain that constitutively expresses RFP (dsRed2.0) under control of the *GRA1* promoter (Oldenhove et al., 2009), we inactivated the endogenous selectable marker *HXGPRT* through transfection with three gRNAs targeting the third, fourth and fifth exons. These gRNA expression vectors were assembled by annealing oligos P1/P2, P3/P4, and P5/P6, ligating into BsaI (NEB) digested pU6-Universal (AddGene #52694), and sequence verifying with P19(Sidik et al., 2014). Transfected parasites were selected with 300 μg/mL 6-thioxanthine and screened for large deletions with P7/P8 (Donald et al., 1996). This strain was made constitutively Cas9^+^ by co-transfection with pCas9-CAT (AddGene #80323) and pU6-Decoy (AddGene #80324) as described previously (Sidik et al., 2016). The strain was further transfected with ScaI (NEB) linearized pBAG1-mNeonGreen, which contains the promoter of BAG1 (1.22 kb upstream of the coding sequence ATG, amplified with primers P9/10) driving expression of mNeonGreen, and also contains a HXGPRT resistance cassette. We selected for integration with 25 μg/ml mycophenolic acid and 50 μg/ml xanthine (Behnke et al., 2008; Bohne et al., 1997). Note this plasmid contains two identical DHFR 3′ UTRs, and care had to be taken to avoid the loss of HXGPRT by recombination during growth in bacteria.

#### *BFD1*^frameshift^

One gRNA was designed targeting the first exon of *BFD1*. Oligos P89/P90 were annealed, Gibson-assembled into pU6-Universal, and sequenced verified with P19, generating plasmid pU6-BFD1-DHFR. Bradyzoite reporter strain parasites were transfected with 50 μg of AseI (NEB) linearized pU6-BFD1-DHFR, and selected with 3 μM pyrimethamine in standard medium the next day. After stabilization of the population, parasites were subcloned into 96-well plates at 3 parasites per well. Clonal strains isolated from single plaques were screened and sequenced for polymorphisms at the targeted site.

#### ME49∆*KU80*

Two gRNAs were designed targeting regions immediately upstream or downstream of the *KU80* coding sequence. Oligos P11/12, P13/14, P15/16, P17/18 were annealed, Gibson-assembled into pU6-Universal, and sequence verified with P19. An early passage ME49 strain was transfected with 25 μg of each plasmid, and immediately subcloned into 96-well plates at 20 or 40 parasites per well to account for loss of viability during transfection. Clonal strains isolated from single plaques were screened for deletion of *KU80* with P20/21, which amplifies a band of ~5.9 kb in wild-type parasites or ~500 bp if *KU80* is excised. A single mixed population was identified from 225 strains tested, and further subcloned to isolate ME49∆*KU80*. Loss of *KU80* was confirmed by complete sequencing of the locus and failure to amplify an internal fragment using P22/P23.

#### ME49∆*KU80*∆*BFD1*

Two gRNAs were designed targeting regions immediately upstream or downstream of *BFD1*. Oligos P24/25 and P26/27 were annealed, Gibson-assembled into pU6-Universal, and sequenced verified with P19. A repair template consisting of the *SAG1* promoter driving expression of mNeonGreen was amplified from p*SAG1*-mNeonGreen using primers P28/29 with 40 bp of homology to regions flanking the targeted sites. ME49∆*KU80* was transfected with 50 μg of each gRNA and 10 μg of repair template. 5 days post-transfection parasites were sorted by green fluorescence and subcloned. Clonal strains isolated from single plaques were further characterized by sequencing the locus using P30/31 to confirm complete deletion of *BFD1*.

### Immunofluorescence assays

HFFs were grown on coverslips for 2–3 days before inoculation with *Toxoplasma*. Coverslips were fixed with 4% formaldehyde for 20 minutes, permeabilized with 1% Triton-X 100 for 8 minutes, and blocked (5% normal goat serum and 5% IFS in phosphate buffered saline (PBS)) for at least 15 minutes (Figs. 1, 2, 4, and **S12**). Alternatively, fixation was done using ice-cold 100% methanol for 2 minutes, without further permeabilization, followed by blocking as above (**Fig. S8**). All primary and secondary antibody incubations were performed for 1 hour, with coverslips inverted on 50 μL of antibody dilutions in blocking buffer on Parafilm in a humidified chamber. Three washes with PBS were performed after each step. Coverslips were mounted on 5 μL of Prolong Diamond (ThermoFisher) and set for 30 minutes at 37°C or overnight at room temperature. DBL-488 (Vector Labs) was used at 1:500. Mouse-anti-Ty antibody (BB2) was used at 1:1000 (Bastin et al., 1996). Rabbit-anti-GAP45 was a gift from Dominique Soldati (University of Geneva) and was used at 1:1000 (Plattner et al., 2008). Rabbit-anti-SAG2Y was used at 1:2000 (Saeij et al., 2008). Mouse-anti-SAG1 (DG52) was used at 1:500 (Burg et al., 1988). Hoechst 33258 (Santa Cruz) was used at 1:2000. Secondary antibodies labeled with Alexa Fluor 488, 594 or 647 (ThermoFisher) were used at 1:1000.

### Quantification of gene disruption

Bradyzoite reporter strain parasites were transfected with 50 μg of AseI (NEB) linearized pU6-SAG1-DHFR (AddGene #80322), encoding a gRNA targeting *SAG1*. Selection with 3 μM pyrimethamine in standard medium was initiated the next day, and drug-resistant pools were inoculated onto coverslips two passages (five days) after transfection. Coverslips were fixed 24 h later with methanol, and stained for SAG1 to quantify KO rates, relative to an untransfected control. GAP45 was used as a counterstain. Knockout rates were quantified before each screen to ensure Cas9 activity.

### Endogenous tagging

To endogenously tag TGME49_312330 and TGME49_208740, ME49∆*KU80* was co-transfected with 50 μg of pCas9-CAT and 50 μg of BsaI-linearized p312330-Ty or p208740-Ty. Selection with 3 μM pyrimethamine in standard medium was initiated the following day. Parasites were subcloned in 96 well plates, and isolated clones screened for successful integration using primers P86/88 or P87/88, respectively, and validated by Sanger sequencing.

### Overexpression vectors of *BFD1*^WT^ and *BFD1*^∆DBD^

The sequence of *BFD1* was amplified from ME49 cDNA using primers P32/33. To amplify *BFD1* lacking the DNA-binding domain (removing amino acids 921–1019), primers P32/34 and P35/ P33 were used. *BFD1* fragments were combined by Gibson Assembly (NEB) with the TUB1 promoter (amplified with P36/37) and the native *BFD1* 3′ UTR (~1.1 kb amplified with P38/39), and verified by Sanger sequencing with oligos P79-P85.

### Phylogenetic analysis of *BFD1*

Protein sequences containing SANT/Myb-like domains were obtained for representative apicomplexan genomes from EupathDB based on their annotation with SMART domain SM00717. Domains from human c-Myb and CDC5L were used for comparison. Individual domains were extracted from each sequence and aligned using ClustalW, and the phylogenetic tree was generated by neighbor-joining (**Fig. S5**)(Larkin et al., 2007). Alignments were also prepared for the concatenated domain sequences for a subset of proteins, and Bootstrap values were calculated for 10,000 trials (Fig. 2A). Visualizations generated using FigTree (v1.4.4).

### Library assembly

The gRNA oligonucleotide library was synthesized by Agilent and resuspended at 1 ng/ μL in water. All library amplifications were done using iProof (Bio-Rad), using 2.5 ng of the oligonucleotide pool as template per 50 μL reaction. Sublibraries were amplified using primers P40/41 for library 1 and P42/43 for library 2, and subsequently amplified with primers P44/45 for cloning. Amplified libraries were Gibson assembled into gel-extracted (Zymo) BsaI-digested pU6_Library_DHFR, dialyzed against water, and electroporated into E. cloni electrocompetent cells (Lucigen) according to manufacturer’s protocol. Coverage was assessed by dilution plating in comparison to a no-insert negative control. Libraries were maxiprepped (Zymo), and retransformed into chemically competent NEB 5-alpha (NEB) to improve yields. Both E. cloni and NEB 5-alpha libraries were sequenced to ensure diversity. Libraries were linearized with AseI, dialyzed 1 h against water, and divided into 50 μg aliquots. Guide RNAs against mNeonGreen were assembled separately by annealing primer pairs P46/47, P48/49, P50/51, P52/53, or P54/55 and Gibson assembling into gel-extracted, BsaI-digested pU6_Library_ DHFR. Constructs were verified by sequencing with P19, and spiked into library aliquots at equimolar concentrations.

### Cas9-mediated genetic screening

Reporter strain parasites were grown up in ~10 15-cm plates per screen. 10 transfections were performed for each library as described previously, with 50 μg of library transfected into approximately 2.6 × 10^7^ parasites in 400 µL cytomix for each reaction (Sidik et al., 2014). Transfections were pooled and split between four 15-cm dishes. Medium was changed the following day to standard media supplemented with 3 μM pyrimethamine and 10 μg/mL DNaseI (Sigma-Aldrich). At each passage of the screen, plates were scraped and parasites were mechanically released with a 27-gauge needle. For the second passage of the screen, all parasites were passed into four 15-cm plates without counting. All subsequent passages were performed at a multiplicity of infection (MOI) of ~1 (6 × 10^6^ parasites per plate). Plates lysed every 2–3 days under unstressed growth in standard media supplemented with 3 µM pyrimethamine. At fourth passage (14 days post-transfection), parasites were inoculated into seven 15-cm plates, and medium was changed after 4 hours to standard medium supplemented with 3 µM pyrimethamine (3 plates) or alkaline stress media (4 plates). Unstressed parasites were passaged at an MOI of 1 every 2–3 days into one or two 15-cm plates, in standard medium supplemented with 3 µM pyrimethamine. Parasites under stressed conditions did not lyse out and were not passaged for the duration of the experiment, and medium was changed every 2 days to fresh alkaline stress medium. At each passage of unstressed parasites, 1–4 × 10^7^ parasites were frozen down. At 10 days post media change, stressed populations were scraped, parasites were mechanically released, passed through a 3 µm filter, and sorted based on green fluorescence. Sorting was done using a BD FACS Aria II. At final timepoints for stressed parasites, both bulk populations (~2 × 10^5^ parasites) and mNG^+^-sorted populations (~7 × 10^5^) were frozen. DNA was isolated using the Qiagen Blood and Tissue kit, following the protocol for blood cells. Integrated gRNAs were amplified and barcoded using primers P56 and P57–76 in 50 μL reactions. Each reaction contained 200 ng or a maximum of 20 μL of template DNA. Amplicons were gel-extracted (Zymo), eluted in water, and quantified using the QuBit dsDNA HS kit (ThermoFisher). Amplicons were pooled equally at a final concentration of 8 pM each and sequenced using a MiSeq v2 kit. Reads were 40 bp single-end and an 8 bp index. Custom sequencing primer P77 and custom indexing primer P78 were used. Guide RNAs were quantified using a custom Perl script. Guides not detected were assigned a pseudocount of 90% of the lowest detected gRNA in that sample. The phenotype or differentiation score for a gene was calculated by determining the mean log_2_-fold change of all five gRNAs targeting that gene in the final sample compared to the input library. All analyses were done in R.

### Stage-specific RNA-sequencing and analysis

Parasites were allowed to invade and replicate inside host cells for 24 h in standard medium, and then switched to either standard or alkaline stress medium. For FACS, parasites were mechanically released from host cells using a 27-followed by a 30-gauge syringe needle, and passed through a 3 µm filter. At 24 and 48 h post medium change, ~1 × 10^5^ unstressed mNG^−^ or stressed mNG^+^ parasites were sorted directly into TRIzol LS and frozen on dry ice. Sorting was done using a BD FACS Aria II, and visualization of events and gates using FCS Express 6. RNA was extracted by TRIzol-chloroform according to manufacturer’s protocol, DNaseI digested, and TRIzol-chloroform extracted again. RNA quality was assessed using a BioAnalyzer or Fragment analyzer. When possible, two samples were prepared per replicate and timepoint and treated as technical replicates in downstream processing. Libraries were generated using the SMARTseq low-input v4 kit, and sequenced on two lanes of a HiSeq 2000. Reads were 75 bp, paired-end. Alignment to the ToxoDB v. 36 assembly of the ME49 genome was done using STAR (Dobin et al., 2013). Differential expression analysis was done using the DESeq2 R (v. 1.21.16) package (Love et al., 2013). The cutoff for differential expression was an adjusted *p* value of 0.001 or lower.

### Single-cell RNA-sequencing and analysis

Seq-Well was performed as previously described, with the following amendments to the protocol. Following Exonuclease I treatment, second strand synthesis was performed. Nextera XT library reactions were done using 1 ng of WTA input, 11 μL TD and 4 μL ATM per reaction. (Gierahn et al., 2017). Single cell suspensions of *Toxoplasma* were prepared by syringe release of parasites from host cells with a 27 followed by a 30-gauge needle, followed by filtering through a 5 µm filter and counting on a haemocytometer. Approximately 12,000 parasites were loaded per array, with two arrays per strain and timepoint used for stressed samples, and one array per strain for unstressed. At the 48 h timepoint, one wild-type stressed and one ∆*BFD1* stressed array failed to seal correctly, resulting in only one array per strain and growth condition at this timepoint. Sequencing was done on two NovaSeq flowcells. Pre-processing, alignment to the ToxoDB v.41 assembly of the ME49 genome, and downstream processing done following the DropSeq Cookbook (http://mccarrolllab.org/dropseq/) (Macosko et al., 2015). An estimate of the number of single cells was made using plotCumulativeFractionOfReads() from Dropbead in R with a maximum of 12,000 cells(Alles et al., 2017). The corresponding cells were then further parsed and analyzed using the Seurat R package (v. 2.3.4) (Butler et al., 2018).

In our analysis of all timepoints, genotypes and growth conditions (Fig. 3E-G), we required cells to contain a minimum of 200 and a maximum of 10,000 non-rRNA mapping unique molecular identifiers (UMIs) and have 40% or fewer total UMIs originating from rRNA. In our analysis of the 72 h timepoint (Fig. 3A-B,H-I), we required cells to contain a minimum of 500 and a maximum of 5,000 non-rRNA mapping UMIs, and have 10% or fewer total UMIs originating from rRNA. These more stringent parameters result from higher confidence at this timepoint of what is and is not a cell, based on the tighter distribution of rRNA content and UMIs, with an upper limit of UMIs corresponding approximately to the mean + 2 standard deviations.

Variable genes were identified through outlier analysis of an average expression/dispersion scatter plot using FindVariableGenes() and the following parameters: mean.function = ExpMean, dispersion.function = LogVMR, x.low.cutoff = 0.0125, y.cutoff = .5. Cells were log-normalized and scaled to 10,000 UMIs, regressing out the number of UMIs detected. Principal component analysis (PCA) was run using these variable genes. The number of principal components (PCs) chosen to use for clustering and UMAP visualization was based on permutation analysis and visual inspection of standard deviations of PCs.

For clustering of 72 h timepoint unstressed parasites only (Fig 3. A-B), principal components 1–6 were identified as statistically significant (*p*-value < .001) by permutation as implemented by Seurat in JackStraw() and used. These PCs agreed well with visual inspection of the elbow seen when plotting the standard deviation of principal components. Clustering was done using shared nearest neighbor (SNN) as implemented by Seurat in FindClusters() using default parameters and a resolution of 0.6. Visualization by UMAP was done using a min_dist parameter of 0.3. For clustering of all parasites, timepoints and genotypes (Fig. 3 E-F), principal components 1–10 were selected for use, based on the elbow seen when plotting the standard deviation of principal components. Clustering was done as above using default parameters and a resolution of 0.8, and visualization by UMAP used min_dist = 0.5. For clustering of 72 h timepoint unstressed and stressed parasites (Fig. 3H-I), principal components 1–18 were identified as statistically significant by JackStraw() and used, and agreed well by elbow plot. Clustering was done as above using default parameters and a resolution of 0.8, and visualization by UMAP used min_dist = 0.5.

Differential gene expression between clusters or groups of clusters was performed using the Wilcoxon rank sum test as implemented by Seurat in FindMarkers(), using the following parameters: only.pos = TRUE, logfc.threshold = 0.5. For integration of the synchronized tachyzoite cell-cycle microarray data, raw microarray values were scaled and mean-centered per probe. For each cluster (Fig. 3A), probes corresponding to differentially expressed genes were subset from the scaled and centered transformation, and averaged for each timepoint. Cluster 4 did not contain any differentially expressed genes and was therefore excluded from analysis. G1, S/M, and bradyzoite-specific gene signatures were derived from existing literature and from the stage-specific RNA-sequencing performed in this study (**Data S3**) (Suvorova et al., 2013). Cell scores calculated in Seurat using AddModuleScore() with default parameters.

## ACKNOWLEDGMENTS

We thank Dominique Soldati-Favre for the GAP45 antibody, Michael W. White for sharing the synchronized tachyzoite microarray data, and Oliver Billker, Rebecca Cottman, Jacob Waldman, Benedikt Markus, and George Bell for helpful discussions and support. Tyler Smith generated the strategy and constructs for endogenous tagging, and Elizabeth Costa and Emily Shortt provided key technical support.

## Author contributions

BSW, JPS, and SL conceptualized project. BSW, DS, and MH performed all experiments. BSW performed all data analysis. BSW and SL wrote the manuscript, with input from all authors.

## Data and materials availability

Data will be uploaded to appropriate repositories following initial review. All reagents will be available from the corresponding author upon request.

