## Supplementary material for "Identification of a master regulator of differentiation in *Toxoplasma*": Figure S1-11

**This section includes:**

Figs. S1 to S11
Captions for Movie S1
Captions for Data S1 to S8

**Other Supplementary Materials for this manuscript include the following:**

Movie S1
Data S1 to S8

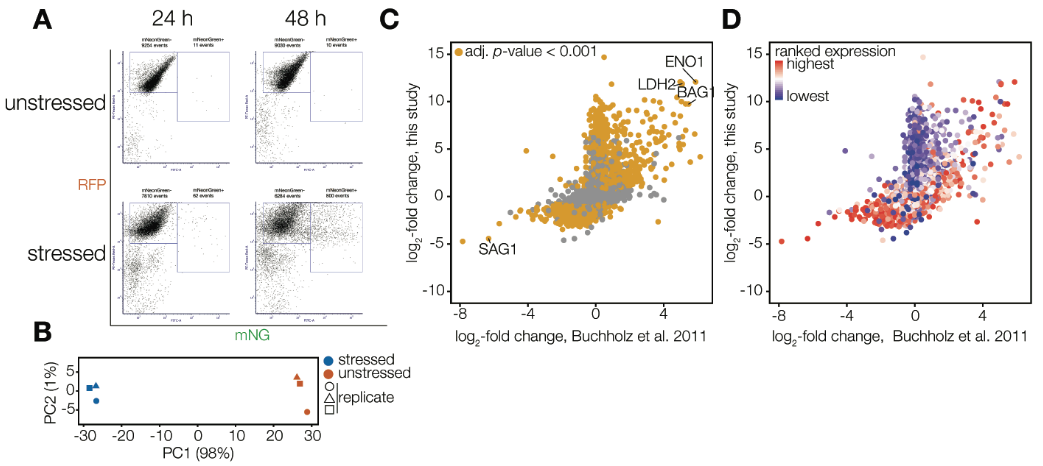


**Figure S1. Stage-specific RNA-sequencing of Toxoplasma.** (**A**) Sample FACS plots of the reporter strain at 24 or 48 h growth under unstressed or stressed conditions. 10,000 events per plot. (**B**) Principal component analysis of stage-specific RNA-sequencing replicates. (**C**) Comparison of differentially expressed genes identified in this study compared to an existing dataset. Color assigned by adjusted *p*-value < 0.001. (**D**) As in C, with color assigned by rank of base mean expression as calculated by DESeq2.

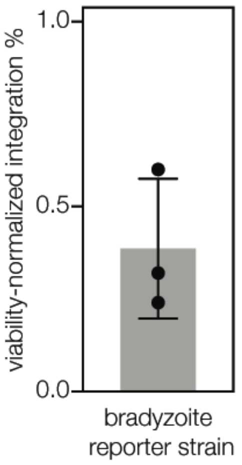


**Figure S2. Integration rates in a non-laboratory-attenuated strain of Toxoplasma are low.** Cas9-expressing ME49 parasites were selected for integration of a gRNA targeting *SAG1*. Plaquing efficiency post-selection was compared to pre-transfection viability rates to obtain a viability normalized integration rate. *n*= 3 independent experiments. Mean ± SD plotted.

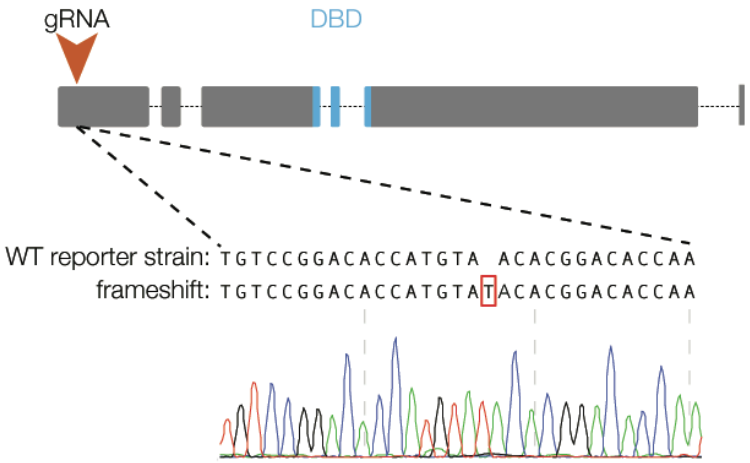


**Figure S3. Generation of a ΔBFD1 reporter strain through Cas9-mediated frameshift.** Transfection of single gRNA targeting the first exon of TGME49_200385 (*BFD1*) into the reporter strain allowed isolation of a strain with a frameshift mutation, resulting in a premature stop codon at amino acid 251.

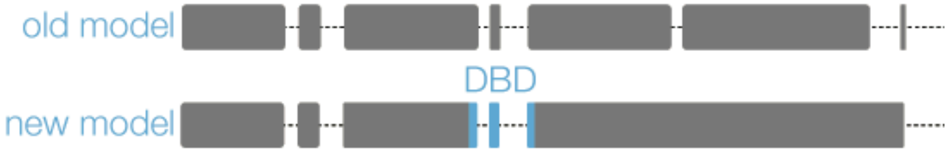


```
>TGME49_200385  (2415 aa)
MEGASTQPMPHSQRERQAPVHAISGTWENCDQAGEYKGTACCVVERQVYSTETHAVEGCCSTRWNG
VKEEKGGGEVSSRTALAGVVHLYTNSDGNAYTHISDQGMRQDEAVAGQEQRHLNCEGEHEKKGNVAA
GGGTLMVTKSEIEPCDDYYSVQRGQSCAGERAPRDGCCRLLDGSRVDPAEGGSEEDENCYTHVLQKH
LGQMPTAPYQGDDRLQTDLRLGDFQSGPSDALSGHHVTRTPILSPHGRYSEYISERLAWQYAERPGHGI
AAENTVVMHSHTGEATGSLRADAPSQWSTESRLQFHVGSQFTTENPELFAGIVGLDTEQFDARNAEALH
WRQQGEARSTETECIADITPERGNRSTIRQWNKPEAARQEYHRASASARGPEETNIITRGAHSDDQSIDA
AAPGCWAARHLTSRQHPNPRPRMKEEHCGREREVLSSEQPSDCGETQKTPASHSLVLDSKSRADASQ
TSPYTRNDEIQTIRAFEEQCLEVQLPQSRLRRGIDYEAGMSQLAPTTQELKAKVLTTGLSPSCPNAAVLRA
SVPLSMDTTVTVHLADVEDPSSKRHTPIENSISSGHCDPPCSSPGGMMEPRVSVSGLGSPHNCVEDREI
ARHGDGIGRDILERRPLPFSPAALLTGCSSTAASSVVSGDVFTSVAASARANACRSSTDTQEESGCGDAP
RYVLLFPDEQDERLGHAQTSIATPSGSGQSKCMFKGDGNAEATFREEVKSSELRTPSRIQTRRLLPGVQ
LQGMDCDGCGASDPQKGRPAPETGFLPEICHFSPQHPWQPGSEVNQGYRAGADYGTSRVQQSLENC
SWEESVDEGEQPRTTSSSSYGQDTQKRDSFLQSIDNRIASEPVDPTGTCGYNSTPETFRSGCIMRDQM
LGVKPHLSHVSLTREGKHRGQQTHLRKTGVPTWSAEEDASLAELVSRKGFKWALISSQLTGAFGIPRTG
KQCRERWFNHVNPEVKKGDWSAEEDAMILMLQNELGNRWATIAKKLRGRTENAVKNRFISLSNARLGY
GRPKRDGSSADCFSNRRTGSGKSSGITGMPNLCQSVCSAGTTKKDSSESGNHFVMSVATKVFEFSDVA
VDSGVSRPRQCTGTSPSCGHPSAGEGDPSHLKNTDVVGREQPIQRNNNESGKAAEQTAFSGVKTGTL
SVSQDAVPVGRLVVASVGPQHIRRSFPTDETLPKFAAKEHNNQQLNDEREHLEQSNSTSEGSFLASAHE
HADIARSDPDEDTLEPHQKRRRKRCAIAYQGEERGDSNGLDSIADRAEQAGNFQAMRKANTDNGKVDY
LEPHRYEKLSPCEQVIQPSLRPACDHRGAPQNSVESGEQSPDAQRQLCNQGCRTSNRTVHSSVYSNE
VESNELRGVFRLAEQSLPSQSGDPAWSTAGFQLSILPQKVEVHSRNKCDGQNVMYRCSPGSLPTTHQQ
TVFHYDRDSSRFPCAAKPAAASGAQGTIEENDGLVKEGPSMIVTGSSVEVVHCCSVSLRRRDRSLPSAQ
LWTSQETESDTNPSPNQQHESCHQYCKRHAAWCGKTDQFSKLTSSHQENSSGKDACLVSVSPTVTLD
DLQKQSRGTVLSAKEEIGKPETWSHVVDNTYSKTDHQRASLCAENSSGCAEGSTELVRFSGGSVKSGS
SMSVDCGNGNPDDCQDCKAEEIWRGEQRYAERGHSVESRGAGSVGRSTDLTITDSGSMPLCASPIGR
PPADNDTLFLSDARCNVVAQLNHQDNSRISRLASCEEEFLACGGERLINASGGFKPDGGCLYRMQQAG
ACNTKLHRPVHSCSTIDSEQLEDLPSVEKAVGDRSFSSKRKGDIPPFAEWKKNDELRELYRGVSEAVSH
GQPGDWNGSWPGISGRAHQTSSCFPDRVNASDRRELNSWRLHVSAAAELGSSHIWNSQSYASASVS
RDKQREPPKNGLTGCDVPEYLGTSQSAGLPAANAHERGNFYGHDRCRPREGERVRWVGLQRNRKPE
ASVSSGASNSATTARPKDSTEPDEGNSEGVSTRRKDSGSTAATISRAVSLGMVTPSAACENSSSLTDTSP
PLSHRPSFSFTHCCEETLSRCNSSNYLCPPATCHTSDDGRSLGPSREAQALRSLSLASGYGYPGIPAEA
TSFWQGSSLEHSIMEPQMVPSDDELRLWVHPRDAANWSQSTLKPVAVVSGTDAGDDQHKTPENLTPE
SGQAHRRDGHDMQRVQRCDDEGECPPTTVELTFPHSHSSDEMQDLPSKVQGNFLLRRELSDSLQHET
AESVAGYGWMRIRNAGDIPNSKVPCAWEQCMPASERERGVNDHMSSEASRMSKAASSSFVPSSCTDA
PVVRVGEDTTKSVCEEQQLCEGGNRGSLSPEATGFESLGPPLQLLLVDGYTPFEPVVEKVSQTMEQTLF
PVPGQETDTRDEDGRYNCECLQNRQPPLHSGGLM
```

**Figure S4. Updated gene model and protein sequence of TGME49_200385.** Sequencing of cDNA suggested the 5th, 6th, and 7th exons as annotated on ToxoDB v. 42 are a single exon, which results in a change of reading frame of the final exon. DNA-binding domains (SM00717) highlighted in blue.

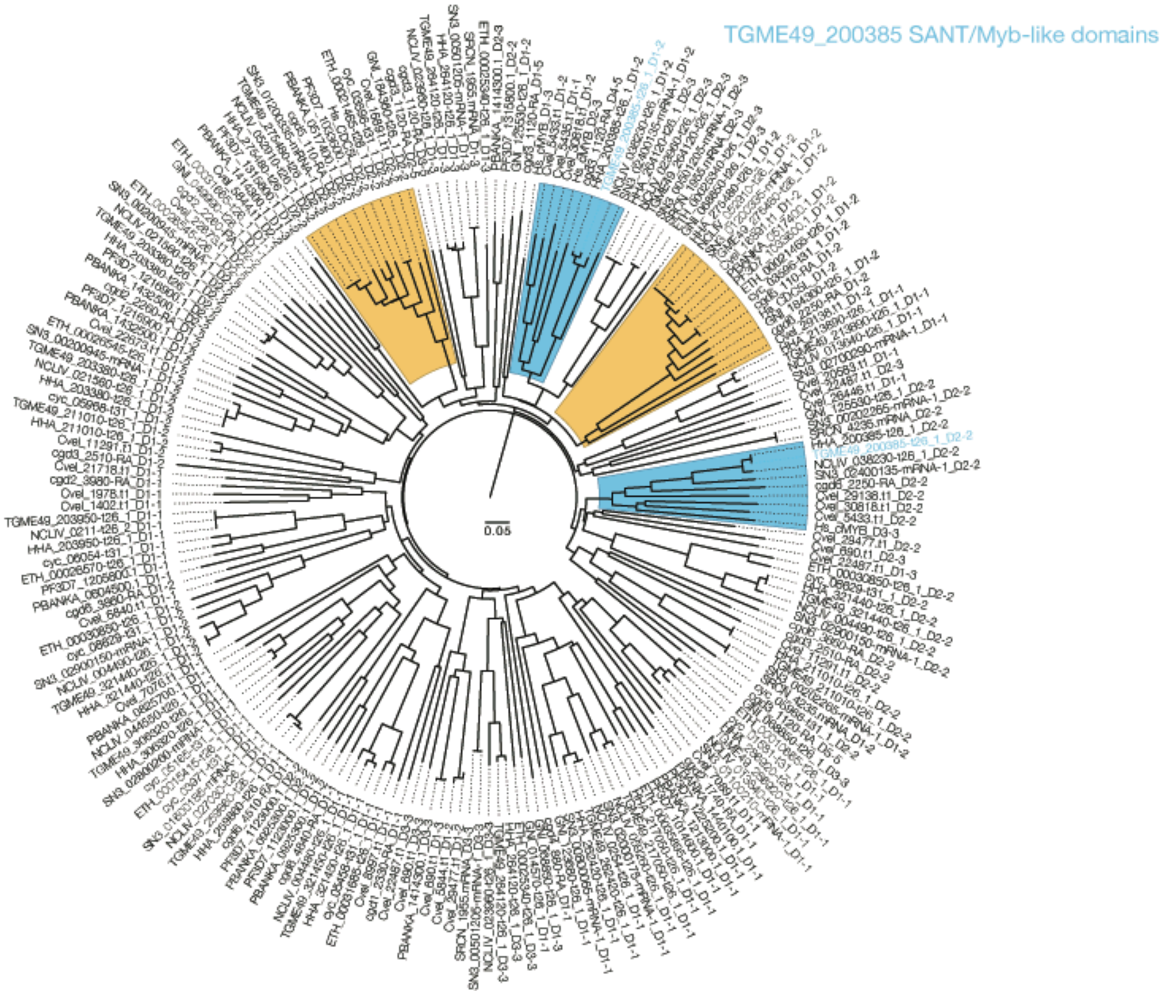


**Figure S5. Phylogenetic analysis of Toxoplasma SANT/Myb-like domains.** Neighbor-joining phylogenetic tree of SANT/Myb-like DNA-binding domains (SM00717) present in representative Apicomplexan genomes, along with human c-Myb and CDC5L. Clades containing c-Myb and CDC5L are highlighted in blue and orange, respectively. Alignment performed using ClustalW. Scale bar is substitutions per site.

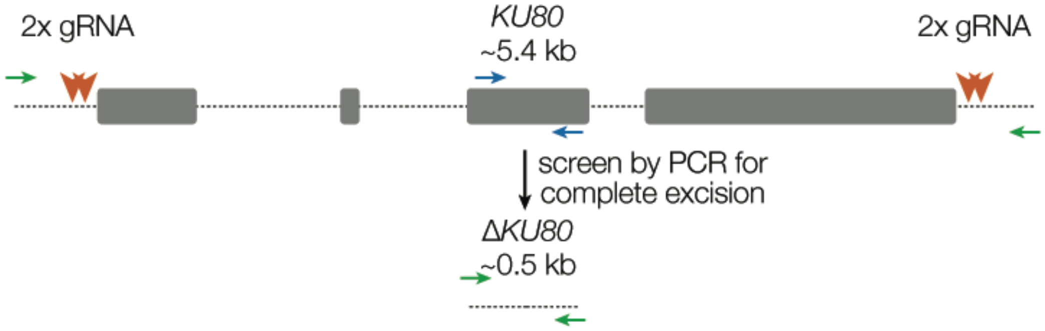


**Figure S6. Strategy used to generate an early passage ME49ΔKU80.** Two gRNAs were designed against both the 5′ and 3′ of the *KU80* locus. Transfection of all four gRNAs into an early passage ME49 strain followed by immediate subcloning allowed recovery of a clone that had deleted the intervening sequence.

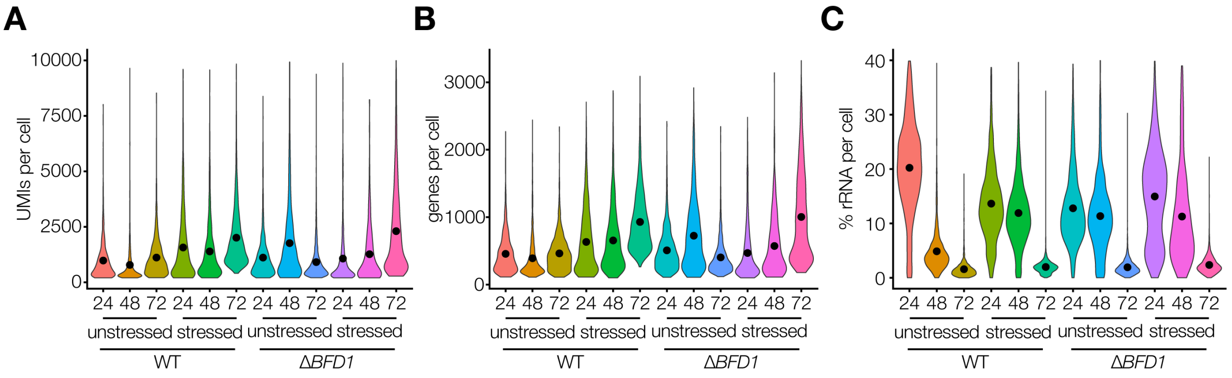


**Figure S7. Gene detection is maximized and rRNA content minimized at the 72 hour timepoint.** (**A**) Distribution of UMIs across single cells from indicated samples and timepoints. Pre-processing quality control cutoffs required a minimum of 200 and maximum of 10,000 UMIs. (**B**) As in A, but unique genes detected. (**C**) Percentage of UMIs corresponding to ribosomal genes. Pre-processing quality control cutoffs allowed a maximum of 40% rRNA reads.

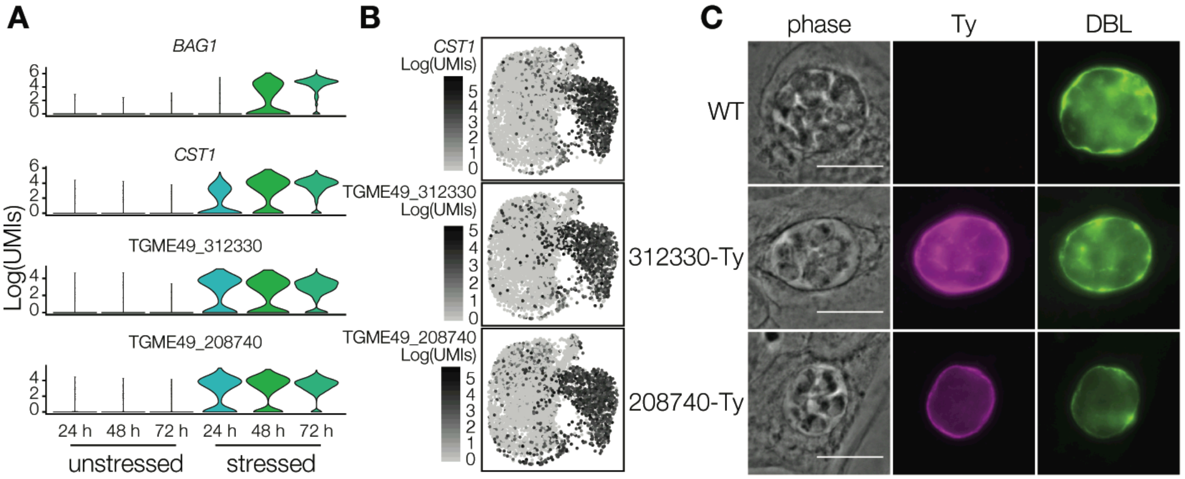


**Figure S8. TGME49_312330 and TGME49_208740 are novel cyst-wall proteins robustly expressed earlier than canonical bradyzoite markers.** (**A**) Violin plots of expression of indicated genes in wild-type parasites after 24, 48 or 72 h growth under unstressed or stressed conditions. (**B**) UMAP visualization as in **Fig. 3E** colored by expression of indicated genes. (**C**) Endogenous tagging of TGME49_312330 and TGME49_208740 localizes both to the cyst wall. Pictures taken 72 h post alkaline stress. Scale bar is 10 μm.

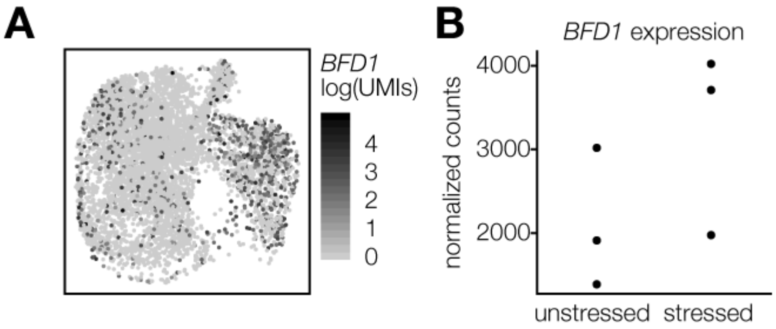


**Figure S9. Expression of BFD1 is not stage-specific, but is upregulated in bradyzoite-containing clusters.** (**A**) UMAP visualization as in Fig. 3E colored by expression of *BFD1*. (**B**) Normalized counts for *BFD1* in stressed or unstressed stage-specific RNA-sequencing replicates.

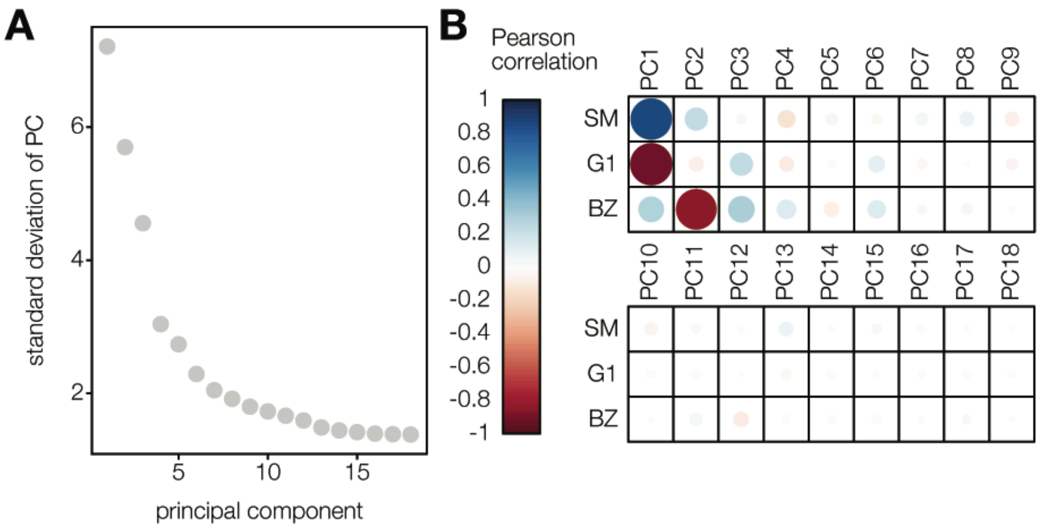


**Figure S10. The majority of variance is driven by cell-cycle and stage-specific genes.** (**A**) Plotting the 18 principal components (PCs) determined to be statistically significant by permutation analysis. The first three PCs explain the majority (66.4%) of variance at the 72 h timepoint. (**B**) Pearson correlations of cell embeddings in PCs 1–18 to cell scores for G1, S/M, or bradyzoite-specific gene signatures.

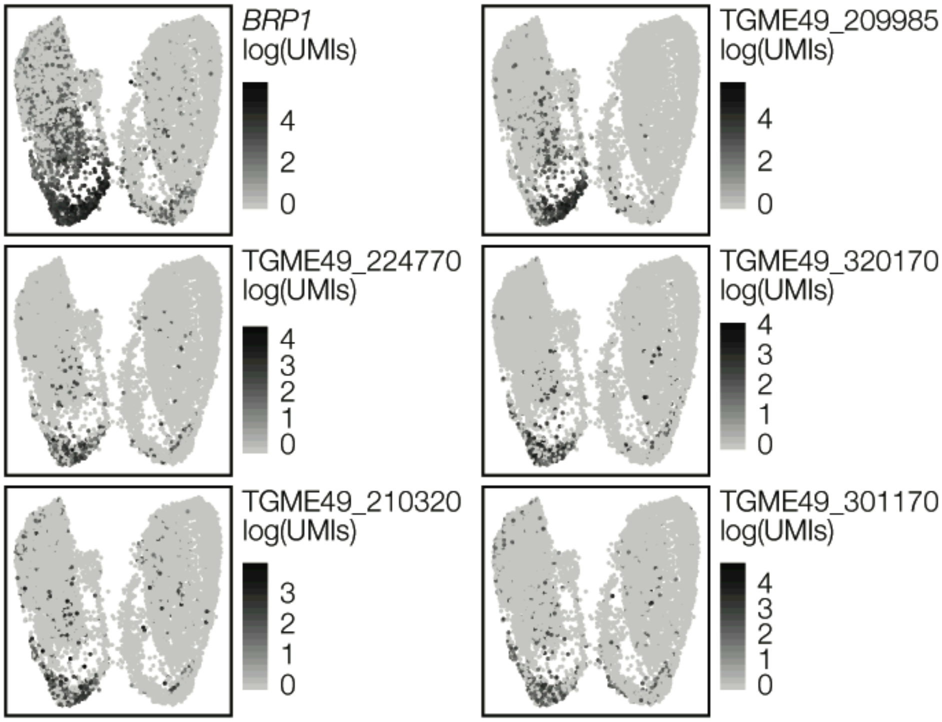


**Figure S11. Single-cell RNA-sequencing identifies markers specific to replicating bradyzoites.** UMAP visualizations colored by expression of indicated genes identified as upregulated in replicating bradyzoites.

**Movie S1. A *BFD1*$^{frameshift}$ reporter strain fails to express mNG under alkaline stress.** Parasites were allowed to invade host cells for 4 h under unstressed conditions, then shifted to alkaline stressed conditions and imaged. One image was taken per hour over 72 hours. RFP is in magenta and mNG is in green. Scale bar is 20 µm.

**Data S1. Differential gene expression analysis of stage-specific RNA-sequencing.** $Log_2$-fold changes correspond to bradyzoites compared to tachyzoites, as shown in **Fig. 1D**.

**Data S2. Data from and libraries used in Cas9-mediated screening for differentiation mutants.** Both raw gRNA counts and analysis at the gRNA and gene level are provided.

**Data S3. Gene modules used for scRNA-seq analysis.**

**Data S4. Differentially expressed markers identified for scRNA-seq clusters corresponding to the tachyzoite cell cycle.** Cluster names and cells used correspond to **Fig. 3A**. Log-fold changes correspond to cells in the named cluster compared to cells in all other clusters.

**Data S5. Differentially expressed genes identified between unstressed and stressed wild-type or Δ*BFD1* parasites at each scRNA-seq timepoint.** Log-fold changes correspond to alkaline-stressed parasites compared to unstressed parasites.

**Data S6. Differentially expressed genes identified between tachyzoite and bradyzoite-containing clusters in scRNA-seq**. Tachyzoite clusters (4, 5, 6, 8 and 9), bradyzoite clusters (0, 1, 7, 11 and 12), and cells used correspond to **Fig. 3H**. Log-fold changes correspond to bradyzoite-containing clusters compared to tachyzoite-containing clusters.

**Data S7. Differentially expressed markers identified for scRNA-seq clusters corresponding to tachyzoites, bradyzoites, and stressed Δ*BFD1* parasites.** Cluster names and cells used correspond to **Fig. 3H**. Log-fold changes correspond to cells in the named cluster compared to cells in all other clusters.

**Data S8. Primers and plasmids used in this study.**
